# Epigenetic reprogramming induced by Acetyl-CoA and SAM depletion is an evolutionarily-ancient path to malignant growth

**DOI:** 10.1101/2024.12.10.627655

**Authors:** Zhe Chen, Xiaomeng Zhang, Mingxi Deng, Chongyang Li, Thi Thuy Nguyen, Min Liu, Kun Dou, Toyotaka Ishibashi, Jiguang Wang, Yan Yan

**Affiliations:** Division of Life Science, The Hong Kong University of Science and Technology, Hong Kong SAR, China; Shenzhen PKU-HKUST Medical Center, Shenzhen, China; Department of Chemical and Biological Engineering, State Key Laboratory of Molecular Neuroscience, The Hong Kong University of Science and Technology, Hong Kong SAR, China; SIAT-HKUST Joint Laboratory of Cell Evolution and Digital Health, HKUST Shenzhen-Hong Kong Collaborative Innovation Research Institute, Futian, Shenzhen, China; School of Life Science and Technology, ShanghaiTech University, Shanghai, China; School of Life Science, Peking University, Beijing, China

## Abstract

Despite the vast diversity of life forms and living histories, it appears that all branches of Metazoa face the challenge of tumor growth. Contrary to human tumors, which take years to form, tumors in short-living species can arise within days without accumulating multiple mutations, raising the question whether the paths to tumorigenesis in diverse species have any commonalities.

In a fly tumor model caused by loss of cell polarity genes, we first identified the rise of a glycolytic cell population over time, resembling the Wartburg effect observed in human tumors. We further identified two key metabolic changes in these fly tumors. First, a systemic depletion of acetyl-CoA leads to a reduction in histone acetylation levels and stochastic silencing of actively-transcribed genes. Second, defects in the methionine cycle cause a systemic depletion of S-Adenosyl methionine, which further reduces histone methylation levels and causes stochastic activation of transposons. Perturbation of the methionine metabolic process strongly inhibits tumor growth. Finally, to understand the evolutionary origin of tumorigenesis, we performed comparative studies of fly and human tumors, and identified human tumors that exhibit metabolic signatures similar to those of fly tumors. We found that human tumors with high metabolic similarity to fly tumors have a lower mutational load, younger patient age, and lower DNA methylation levels. This study suggests that tumorigenesis processes have a deep evolutionary origin and highlights that depletion of key metabolites is an evolutionarily-ancient driving force for tumorigenesis.

## Introduction

Cancer is not a disease that only affects human. The appearance of tumors seems to coincide with the origin of metazoan. Various tumor types have been documented from hydra, clams worms, flies, fish to mammals^1,2^. Cancers have even been identified in dinosaur fossil records^3–5^. Tumors in *Drosophila Melanogaster* were first reported in 1918^6^, and they represent one of the best-understood tumor models in species outside mammals^7^. For example, three conserved cell polarity genes, *scribble (scrib)*, *lethal giant larvae (lgl)*, and *discs-large (dlg),* are collectively named as neoplastic tumor suppressor genes (nTSGs) ^8–11^. When fly larvae are homozygous mutant for *scrib*, *lgl*, or *dlg*, their imaginal discs and optical lobes, which are proliferative epithelial organs in larvae, grow into amorphous masses and kill the hosts within days^8,12^. Even for well-studied fly tumors, it is unclear how closely they resemble human tumors, because these fly tumors reach immortality within days while carrying only one mutation. In contrast, human cancers typically develop over years and acquire multiple mutations to enter the path of carcinogenesis.

Cancer cells are generally thought as being in metabolic states distinct from normal cells^13,14^. For example, most cells can use glucose as an energy source through conserved steps of the oxidation of glucose to pyruvate, the conversion of pyruvate to acetyl groups and the further oxidation of acetyl groups through citric acid cycle^15^. Warburg showed that tumor tissue slices produced lactate from glucose even in the presence of ample oxygen, which is now known as a metabolic signature of cancer cells (the Warburg effect) ^13,16,17^. Genomics and functional studies have identified a number of metabolic enzymes as frequently mutated to support tumor growth in human. For example, mutations in isocitrate dehydrogenase genes (*IDH1* and *IDH2*) are frequently observed in gliomas and acute myeloid leukemia^18,19^. *IDH1* and *IDH2* mutations lead to the accumulation of 2-hydroxygulatrate (2-HG), which inhibits α-KG-dependent dioxygenases and alters the histone and DNA methylation pattern^20–23^. Mutations in fumarate hydratase (*FH*) and succinate dehydrogenase (*SDH*), which lead to accumulation of fumarate and succinate, predispose patients to various cancer types^24–27^. The accumulation of fumarate and succinate can suppress DNA repair pathway^28,29^ and inhibit PTEN through succination^30^. Phosphoglycerate dehydrogenase (*PHGDH*) is found to be frequently amplified in breast cancer and melanoma samples and increases serine flux to support tumor growth^31–33^. Metabolic processes are deeply conserved across species, and it is unknown whether metabolic changes underlying tumorigenesis in different metazoan species share any commonalities.

In this study, we first noticed that a population of Ldh+ cells emerges in fly tumors over time, resembling the Warburg effect observed in mammalian tumors. We further found that fly tumors exhibit major changes in the glycolysis, TCA cycle and oxidative phosphorylation processes, which lead to a systemic depletion of acetyl-CoA. Consequently, we observed a reduction of H3K9ac and H3K27ac levels and stochastic silencing of actively-transcribed genes in these tumors. Moreover, these fly tumors also exhibit defects in the methionine metabolism process, which leads to a systemic depletion of methyl donor S-Adenosyl methionine, a reduction of histone methylation level, and a de-repression of transposons. We then perform comparative studies between fly and human tumors using metabolic signatures, and identified that human tumors with high similarity to fly tumors have a lower mutational load, younger patient age, and lower DNA methylation levels.

## Results

### A glycolytic Ldh+ cell population arises over time in fly tumors

The fly tumor models caused by the loss of conserved cell polarity genes, including *scrib*, *dlg* and *lgl*, are among the best-understood tumor models outside of mammals^9–11^. Note that the *scrib* and *dlg* mutant are phenotypically identical, as these two proteins function as a complex^10,11^. We used the *scrib* and *dlg* mutant interchangeably in experiments, and the choice of which mutant to use was based on the ease of genetic combination. We have previously used single-cell transcriptomics to profile the fly *scrib* mutant wing disc tumors as they grew from 4 days to 14 days^34,35^. From this dataset, we noticed that a population of Ldh+ cells emerge in the late-stage *scrib* mutant tumors (Figure 1A). We further validated the existence of this Ldh+ cell population through examining the expression of a Ldh-GFP enhancer trap line^36^ in the *dlg* mutant (Figure 1B-C). ^37^Ldh+ cell populations were previously shown to be induced in fly wing disc tumors caused by active Pvr^36^ or Hipk^38^ signals. Similar to what have been shown in these contexts, the rise of Ldh+ cell population in the *scrib/dlg* mutant tumors is also regulated by HIF-1 and PI3K/Akt signals (Supplemental Figure 1). Specifically, depletion of HIF-1through RNAi blocks the rise of Ldh+ cells (Supplemental Figure 1A), and activation of PI3K/Akt signaling through *Pten* RNAi promotes the expansion of Ldh+ cells (Supplemental Figure 1B).

**Figure 1.**
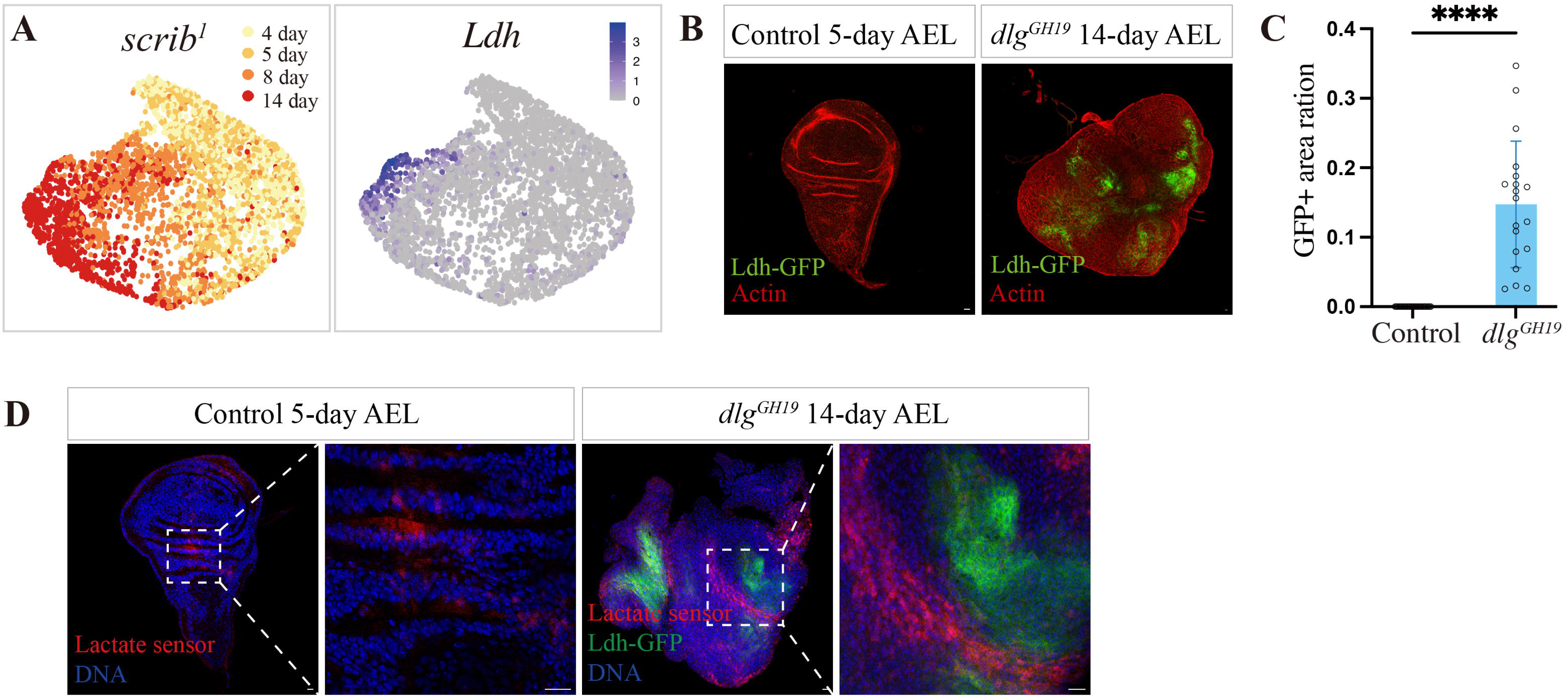
A Ldh+ cell population arises in the fly *scrib* and *dlg* mutant tumors over time. (A) UMAP plot of the *scrib* mutant wing disc single cells from different times and the expression of *Ldh* gene. (B) Control and the *dlg* mutant imaginal discs expressing a Ldh-GFP enhancer trap reporter. Scale bar: 10μm. (C) Quantification of Ldh+ cell percentage in the control and the *dlg* mutant imaginal discs. For B and C, Control genotype: *FRT19A; c855aGal4 Ldh-GFP*, 5-day AEL, n=25. Experimental group genotype: *dlg^GH19^*; *c855aGal4 Ldh-GFP*, 14-day AEL, n=19. (D) Control and the *dlg* mutant imaginal discs expressing R-iLACCO1, a lactate sensor. Control genotype: *c855aGal4/R-iLACCO1*, 5-day AEL, 24/31, n=31. Experimental group genotype: *dlg^GH19^*; *c855aGal4 Ldh-GFP/R-iLACCO1*, 14-day AEL, 21/21, n=21. Scale bar: 10μm.

We further examined whether the expansion of a *Ldh+* glycolytic cell population led to the elevation of lactate levels in the *dlg* mutant tumors using a biosensor for L-lactate, R-iLACCO1^39^. In the control wild-type wing discs, we observed weak fluorescence level at the hinge region (Figure 1D). In the 14-day *dlg* mutant tumors, we observed bright red fluorescence signals (Figure 1D). Notably, the R-iLACCO1 signal is frequently observed in cells adjacent to the Ldh-GFP+ cells and does not completely overlap with Ldh-GFP signals, indicative of possible lactate transfer among these cells^40^. Together, these data suggest that the *scrib* and *dlg* mutant tumors exhibit characteristics of the Warburg effect as these tumors grow over time.

### Depletion of acetyl-CoA led to a reduction in H3K9ac and H3K27ac levels and stochastic silencing of actively-transcribed genes in fly tumors

Glycolysis, the citric acid cycle (TCA cycle) and oxidative phosphorylation are tightly-coupled processes for glucose breakdown and ATP production (Figure 2A). In addition to the rise of Ldh+ cells in these tumors, we further noticed that genes responsible for the breakdown of glucose to pyruvate are collectively upregulated over time in the *scrib* mutant tumors (Figure 2A) (Supplemental Figure 2A). The pyruvate dehydrogenase complex genes, which convert pyruvate to acetyl-CoA for entry into TCA cycle, are collectively downregulated over time in the *scrib* mutant tumors (Figure 2A) (Supplemental Figure 2B). Concurrently, TCA cycle genes are downregulated over time in the *scrib* mutant tumors (Figure 2A) (Supplemental Figure 2C). Moreover, genes encoding components of mitochondrial respiratory chain complexes I-V are collectively downregulated over time in the *scrib* mutant tumors (Figure 2A) (Supplemental Figure 2D). These data suggest that the breakdown of glucose through glycolysis, the TCA cycle and oxidative phosphorylation to generate ATP likely slows down at the step of pyruvate-to-acetyl-CoA conversion for entry into TCA cycle in the late-stage *scrib* mutant tumors.

**Figure 2.**
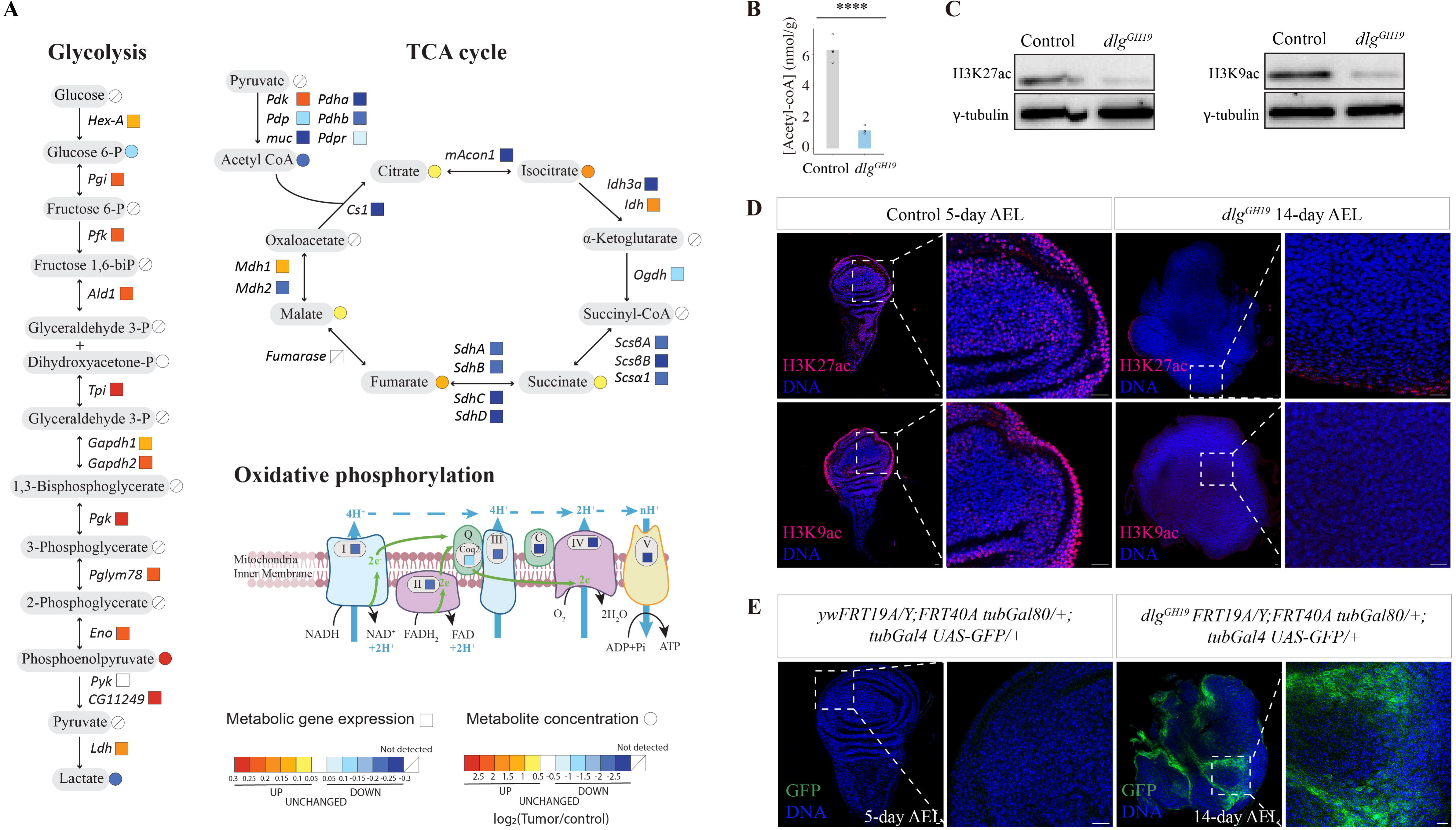
Systemic depletion of acetyl-CoA leads to a reduction in histone acetylation levels and stochastic silencing of actively-transcribed genes in the *dlg* mutant tumors. (A) Plot of metabolic gene expression changes from the *scrib* mutant tumor time-series transcriptomics data (color-coded in squares) and metabolite concentration changes between the 13-day *dlg* mutant larvae and control larvae (color-coded in circles). The gene expression change slope value and metabolite concentration fold change value are color-coded. (B) Plot of Acetyl-CoA concentration in the control and the *dlg* mutant larvae. Control group genotype: *FRT19A,* 5-day AEL. Experimental group genotype: *dlg^GH19^*, 13-day AEL. 4 repeats per group. (C) Western blot of H3K27ac and H3K9ac from the control imaginal discs and the *dlg* mutant tumors. (D) Control and the *dlg* mutant imaginal discs stained for H3K27ac and H3K9ac. Control genotype: FRT19A, 5-day AEL. Experimental group genotype: *dlg^GH19^*, 14-day AEL. H3K27ac, control, n=18, *dlg^GH19^*, n=20; H3K9ac, control, n=22, *dlg^GH19^*, n=15; Scale bar: 10μm. (E) Control and the *dlg* mutant imaginal discs expressing a tubGal80; tubGal4 UAS-GFP reporter. Detection of GFP+ cells is indicative of loss of tubGal80 expression. Control genotype: *yw*FRT19A/Y; FRT40A tub-Gal80/+; tub-Gal4 UAS-GFP/+; n=21, 5-day AEL. Experimental group genotype: *dlg^GH19^*FRT19A/Y; FRT40A tub-Gal80/+; tub-Gal4 UAS-GFP/+; n=33, 32/33, 14-day AEL. Scale bar: 10μm.

We then compared the concentrations of metabolites between the *dlg* mutant larvae and the control larvae through metabolomics profiling (Figure 2A) (Supplemental Table 1). We observe elevated levels of metabolites in TCA cycle including citrate, cis-aconitate, isocitrate, succinate, fumarate, and malate in the *dlg* mutant larvae (Figure 2A), likely indicative of an overall slowdown of metabolite flow through TCA cycle. Notably, in human cancers caused by mutations in *IDH1*, *FH*, or *SDH*, the accumulation of metabolites in TCA cycle, such as succinate and fumarate, is also frequently observed ^20–22,29,30^. In fly brain tumors caused by the loss of Brat protein, the accumulation of TCA intermediates was also observed ^41^, highlighting that the accumulation of TCA intermediates might represent a highly-conserved feature of tumors from different animal species.

Consistent with the downregulation of the expression levels of PDH complex genes, which function to convert pyruvate to acetyl-CoA for entry into TCA cycle, we observed a reduction of acetyl-CoA level in the *dlg* mutant larvae (Figure 2B). Acetyl-CoA not only serves as a metabolic intermediate for TCA cycle and lipid synthesis but also acts as the acetyl donor for protein acetylation, particularly histone acetylation^42–44^. We then examined whether the depletion of acetyl-CoA in the *dlg* mutant larvae affects protein acetylation levels, in particular, histone acetylation levels. Histone PTMs H3K27ac and H3K9ac are classic markers for actively-transcribed chromatin^45–48^. In the *dlg* mutant wing disc tumors, we observed that H3K27ac and H3K9ac levels are much lower than that of the wild type wing discs (Figure 2C-D). Removal of histone acetylation markers can potentially lead to the silencing of actively-transcribed genes. To examine the consequences of an overall reduction in histone acetylation levels in the *dlg* mutant tumors, we adopted a *tubGal80-tubGal4* reporter system which was originally used to monitor loss of heterozygosity in aging intestinal stem cells^49^. In this system, the inactivation of a *tubulin* promoter-driven Gal80 allows the expression of GFP activated by *tubulin* promoter-driven Gal4^49^. The inactivation of tubGal80 can result from either DNA mutations or deletions, or from epigenetic silencing in the *tubGal80* locus^49^. Using this system, we found that GFP+ cells were never observed in the control imaginal discs, indicating that the *tubGal80-tubGal4* reporter system is not leaky (Figure 2E). On the contrary, we frequently observed GFP+ clones in the 14-day *dlg* mutant tumors (Figure 2E). We have previously performed whole-genome sequencing of the *dlg* mutant tumors and did not detected further mutations except the original *dlg* mutation. Therefore, the stochastic loss of Gal80 likely results from the epigenetic silencing of the *tubGal80* locus in association with a loss of histone acetylation markers.

### Depletion of S-Adenosyl methionine leads to a reduction in H3K9me3 levels and transposon activation in fly tumors

We next examined additional metabolic changes observed in the *dlg* mutant larvae in comparison with the wild type larvae, and noticed a significant reduction of methionine cycle-related metabolites such as S-Adenosyl methionine in the *dlg* mutant larvae (Figure 3A-B). S-Adenosyl methionine is a major methyl donor, and therefore we examined whether histone methylation status is affected in the *dlg* mutant tumors. We found that the level of H3K9me3, which is a well-recognized repressive chromatin marker^47,48^, is significantly reduced in the *dlg* mutant tumors in comparison with control (Figure 3C-D).

**Figure 3.**
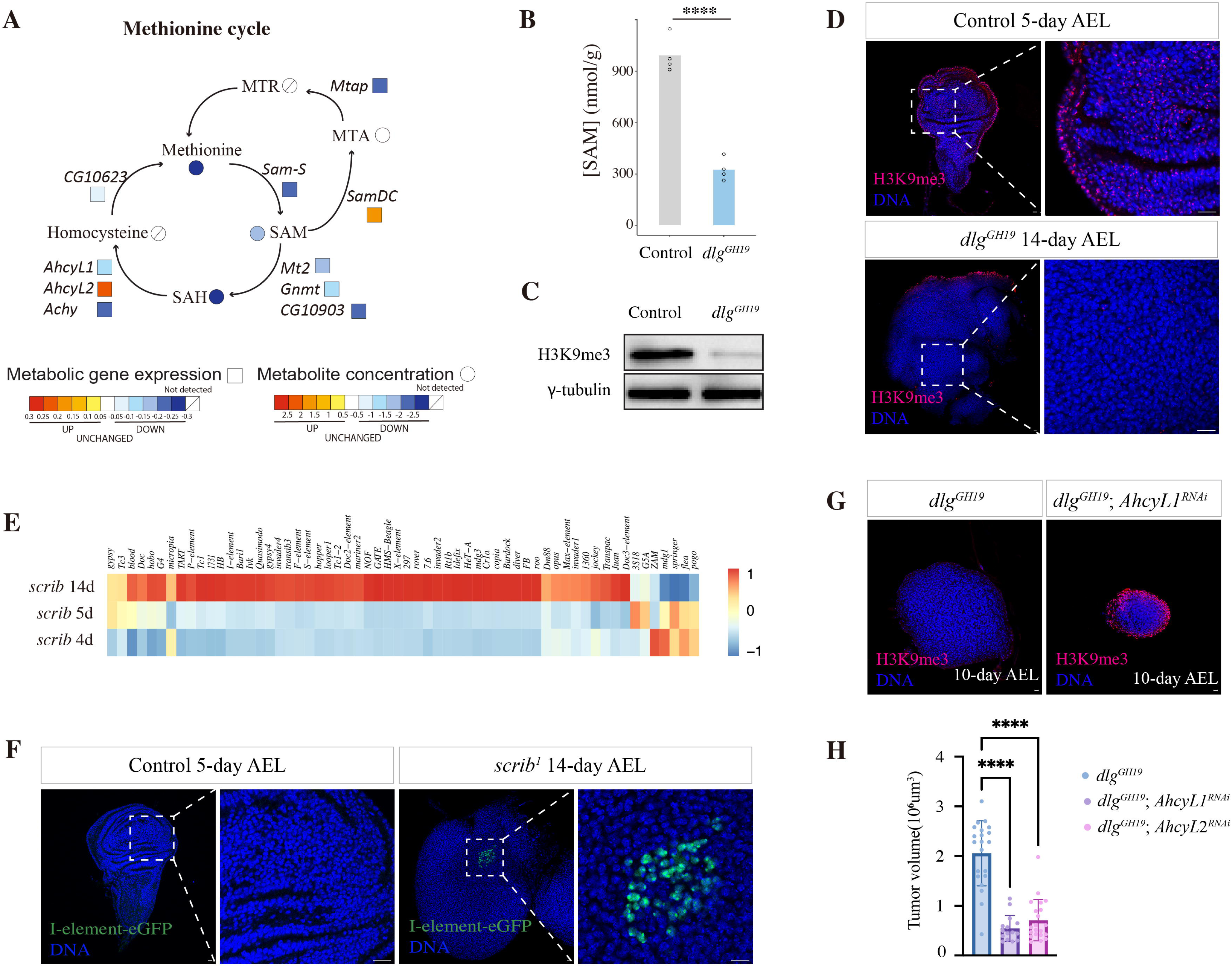
Systemic depletion of S-Adenosyl methionine leads to a reduction in H3K9me3 levels and transposon activation in fly tumors. (A) Plot of gene expression changes from the *scrib* mutant tumor time-series transcriptomics data (color-coded in squares) and metabolite concentration changes between the 13-day *dlg* mutant larvae and control larvae (color-coded in circles) in the methionine cycle. (B) Plot of S-Adenosyl methionine concentration in the control and the *dlg* mutant larvae. Control group genotype: *FRT19A,* 5-day AEL. Experimental group genotype: *dlg^GH19^*, 13-day AEL. 4 repeats per group. (C) Western blot of H3K9me3 from control and the *dlg* mutant imaginal discs. (D) Control and the *dlg* mutant imaginal discs stained for H3K9me3. Control genotype: FRT19A, 5-day AEL, n=21. Experimental group genotype: *dlg^GH19^*, 14-day AEL, n=16. Scale bar: 10μm. (E) Heatmap plot of transposon expression data from the 4-day, 5-day and 14-day AEL *scrib* mutant wing imaginal discs. (F) Control and the *scrib* mutant imaginal discs expressing an I-element-eGFP reporter. Control genotype: I-element-eGFP; FRT82B; n=23, 5-day AEL. Experimental group genotype: I-element-eGFP; *scrib*^1^ FRT82B; n=29, 13/29, 14-day AEL. Scale bar: 10μm. (G) Imaginal discs stained for H3K9me3 (red) and DNA (blue). Scale bar: 10μm. (H) Quantification of tumor sizes for the *dlg* mutant tumors in the control, *AhcyL1 ^RNAi^* and *AhcyL2 ^RNAi^* groups. For (G-H), control genotype: *dlg^GH19^*; *c855aGal4/+*, 10-day AEL, n=21. Experimental group genotype: *dlg^GH19^*; *c855aGal4/AhcyL1^RNAi^;* 10-day AEL, n=15. *dlg^GH19^*; *AhcyL2^RNAi^/+; c855aGal4/+;* 10-day AEL, n=21.

Removal of silencing markers such as H3K9me3 can potentially lead to transposon activation and loss of gene silencing at the heterochromatin region^50^. We examined the transcript levels of multiple transposons in the *scrib* mutant tumors and observed a de-repression of many transposons in the 14-day *scrib* mutant tumors (Supplemental Table 2) (Figure 3E). We then adopted a reporter for transposon activation, I-element-eGFP reporter, which only produces GFP signal upon retrotransposition^51^. In the control wing imaginal discs, we never observed any GFP signal, indicative of tight repression of transposons in the wild type somatic cells (Figure 3F). In the 14-day *scrib* mutant imaginal discs, we observed GFP+ cells in about 50% of samples we examined (Figure 3F), indicating that the reporter was not only reverse-transcribed but its cDNA often successfully landed back into the genome^51^. Together, these results suggest that the late *scrib/dlg* mutant tumors likely harbor a diverse cell population featuring stochastic transposon activation. To determine if depletion of SAM is a driving force for tumorigenesis, we examined the functional consequences of perturbing enzymes in methionine cycle in the *dlg* mutant tumors. Interestingly, we found that knockdown of *AhcyL1* and *AhcyL2*, which potentially elevates SAH hydrolase Ahcy activity^52^, elevates H3K9me3 level and strongly inhibits the *dlg* mutant tumor growth (Figure 3G-H). Note that perturbation of AhcyL1 and AhcyL2 activity in normal wing discs do not affect normal developmental growth (Supplemental Figure 3).

### Identification of an evolutionarily-ancient category of human tumor samples that metabolically resemble fly tumors

Major metabolic pathways are highly conserved across animal species. Using KEGG and Metabolic Atlas as references, we annotated 72 metabolic pathways conserved between fly and human (Supplemental Table 3). We then use transcriptomics data from the *scrib* mutant tumor samples^34^ and 10501 human tumor samples in TCGA database to perform single-sample gene-set enrichment analysis (ssGSEA) for the 72 conserved metabolic pathways (Figure 4A). Using metabolic pathway enrichment scores as a matrix, we further performed correlation analysis to identify human tumor samples with high metabolic similarity to fly tumors (Figure 4A). For the top 7% human tumors with high metabolic similarity to fly tumors (Figure 4B-C) (n=765/10501), they have a younger patient age (Figure 4D), a lower mutational load (Figure 4E), and a lower level of DNA methylation (Figure 4F) in comparison with tumor samples we classified as non-similar to fly tumors. The human tumor samples that metabolically resemble fly tumors show a strong enrichment for mutations in *IDH1*, *GTF2I* and *ATRX* (Figure 4G). GTF2I encodes a transcriptional factor with high mutation frequency observed in thymoma^53^. ATRX is a chromatin remodeler with high mutation frequency in glioma^54^. Consistently, the tumor samples with high similarity to fly tumors are highly enriched in 8 cancer types out of the total 33 cancer types, including B-cell lymphoma, glioblastoma, glioma, ovarian adenocarcinoma, testicular germ cell tumor, thymoma, and uterine carcinosarcoma (Supplemental Figure 4). For the 8 cancer types with a high percentage of tumors metabolically similar to fly tumors, the patient survival curves between the two groups do not show significant differences, except for glioma, where patients with tumors metabolically similar to fly tumors have better survival outcomes (Supplemental Figure 5).

**Figure 4.**
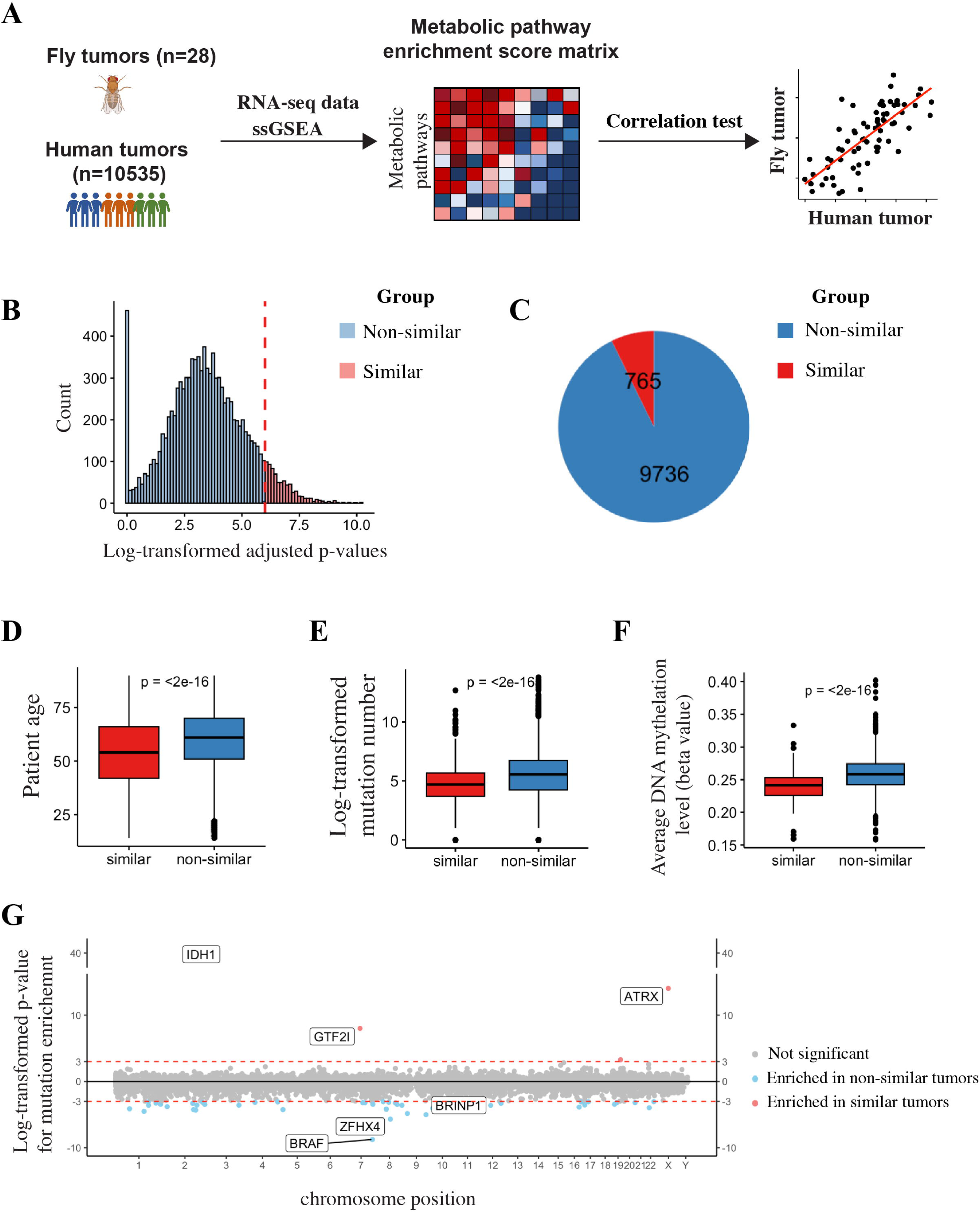
The metabolic gene expression signatures of fly and human tumors. (A) Schematic view of using gene expression data from fly and human tumor samples to identify human tumor samples that metabolically resemble fly tumors. (B) Plot of the distribution of adjusted p-values for metabolic similarity tests between human tumor samples and the fly *scrib* mutant tumors. (C) 765 out of 10501 human tumor samples exhibit metabolic signatures similar to those of fly tumors (D) Patients with tumors metabolically similar to the fly tumors are significantly younger than others. (E) Human tumor samples metabolically similar to the fly tumors acquire a significant lower number of mutations than other samples. (F) Human tumor samples metabolically similar to the fly tumors show a lower level of DNA methylation than other samples. (G) Mutation landscape of human tumor samples metabolically similar to the fly tumors. Dots in the upper panel represent mutation events which are significantly enriched in the metabolically-similar tumor group, while dots in the lower panel are events enriched in the non-similar tumor group.

To further validate these results, we obtained an independent cohort of 185 CGGA glioma tumor data and separate them into two groups based on their metabolic signature similarity to fly tumors (Supplemental Figure 6A-B). Similarly, for this independent dataset, the ages of patients in the similar group are significantly younger than those in the non-similar group (Supplemental Figure 6C). The human glioma samples that metabolically resemble fly tumors harbor a significantly lower number of mutations (Supplemental Figure 6D), show a strong enrichment for *IDH1* mutations (Supplemental Figure 6E), and the patients in the similar group have a better survival outcome (Supplemental Figure 6F).

Together, these data suggest that human tumor samples sharing similar metabolic signatures to fly tumors are less dependent on mutation accumulation over time.

## Discussion

Here we demonstrated that epigenetic reprogramming induced by key metabolite depletion is an evolutionarily-ancient driving force for immortal growth, lowering the barrier to tumorigenesis and the need for mutation accumulation over time. Interestingly, a recent study also demonstrates that the transient silencing of Polycomb complex is sufficient to drive malignant tumor formation in *Drosophila*^55^. Notably, even in human cancers, around 5.3% of tumors do not harbor any known cancer driver mutations^56^. For example, mutations are rarely found in childhood brain tumors^57^. Therefore, epigenetic changes induced by metabolic stress may represent an evolutionarily-ancient route to malignant growth, which co-exist with the acquisition of multiple mutations in long-living species. For the portion of evolutionarily-ancient human tumors, it will be interesting to use fly tumors as a model to test therapeutic interventions. For example, we have found that inhibition of AhcyL1 and AhcyL2, enzymes functioning in the methionine metabolic process, potently inhibits fly tumor growth. In the future, it would be interesting to test if methionine metabolic intervention benefits human patients with evolutionarily-ancient tumors.

The metabolic network is largely conserved among animal species. The accumulation of TCA intermediates is observed in both human and fly tumors. While many studies have focused on the accumulation of oncometabolites^58^, our work here highlight the dire consequences of metabolite depletion, including acetyl-CoA and S-Adenosyl methionine, which are key substrates for histone post-translational modifications. Similar to the concept of oncogenes and tumor suppressor genes, the depletion of metabolites is likely as important as the accumulation of oncometabolites in tumorigenesis processes. A global reduction of histone acetylation and methylation levels can lead to stochastic gene silencing and activation, the implications of which need to be further explored in human tumors.

## Materials and methods

### Drosophila genetics and stocks

*Drosophila Melanogaster* stocks were maintained at room temperature and fly crosses were raised in a 25°C fly incubator. The fly strains used in this study were: *scrib*^1^ FRT82B/TM6B^10^, *dlg^GH19^* FRT19A/FM7C^34^, c885a-Gal4 (BL6990), *Ldh-GFP*^36^, R-iLACCO1^39^, *I-element-*eGFP-reporter^51^, FRT40A tubGal80; tubGal4 UAS-GFP (A gift from Xi lab), *HIF-1* RNAi (BL26207), *Pten* RNAi (VDRC101475), *AhcyL1* RNAi (BL28523)*, AhcyL2* RNAi (BL 61913) and control strains OreR or w1118.

### Wing discs dissection and immunostaining

Wing discs and tumors were dissected, fixed and stained following standard protocols. The primary antibodies used in the study were: rabbit anti-H3K9ac (Abcam Cat No. ab4441, 1:1000), rabbit anti-H3K27ac (Abcam Cat No. ab4729, 1:1000), and rabbit anti-H3K9me3 (Abcam Cat No. ab8898, 1:1000) in 1:1000. The secondary antibodies used were: Goat anti-Rabbit IgG (H+L) Secondary Antibody, Alexa Fluor™ 546 (Invitrogen A11035, 1:1000) and Goat anti-Mouse IgG (H+L) Secondary Antibody, Alexa Fluor™ 546 (Invitrogen A11030, 1:1000). Hoechst 33342 (Invitrogen H3570, 1:10000) and Alexa Fluor™ 647 Phalloidin (Invitrogen A22287,1:1000) were used to stain DNA and actin for cell outline, respectively.

### Western blotting

40 pairs of wing imaginal discs or tumors were dissected in PBS. Tissue samples were vortexed in RIPA lysis buffer for 1 minute, followed by sonication. The sonication program settings included an amplitude of 50, a processing time of 15 minutes, a pulse-on time of 15 seconds, and a pulse-off time of 10 seconds. After cell lysis, the samples were centrifuged at maximum speed for 10 minutes. The supernatant was then measured for protein concentration using the Pierce BCA Protein Assay Kit. The primary antibodies used were rabbit anti-H3K9ac (1:1000, Abcam ab4441), rabbit anti-H3K9me3 (1:1000, Abcam ab8898), rabbit anti-H3K27ac (1:1000, Abcam ab4729), and gamma-tubulin (1:1000, Abcam).

### Confocal imaging and data analysis

Images are acquired on a Leica TCS SP8 confocal microscope. Wing discs and tumors were taking as z-stacks for volume quantification using Image J and CellProfiler^59^.

### Metabolomics

Embryos within a 3-hour window were collected from apple juice plates and aged to larvae of appropriate ages. Wild type larvae (5-day AEL) and the *dlg* mutant larvae (13-day AEL) (12 larvae for each technical replicate, n=4 for each genotype) were sent as frozen samples to Biotree for 600MRM metabolite analysis with LC-MS. Metabolite concentrations, C_M_ (nmol/g), were calculated from the formula below.

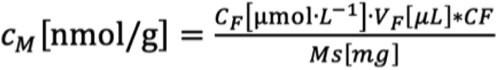

Specifically, C_F_ denotes final concentration, a product of calculated concentration (μmol/L) detected by machine and dilution factor. V_F_ (uL) represents volume of sample. CF is the dilution factor in sample preparation. M_s_ (mg) is the mass of sample. Statistical tests between two genotypes were conducted using t-test.

### Metabolic signature analysis based on transcriptomics data

Bulk and single-cell RNA sequencing data of the *scrib* mutant tumors were obtained from our previous publications^34,35^ (GEO accession Number: GSE130243 and GSE130566). The gene expression data for 10501 tumors of 33 cancer types were obtained from TCGA Data Portal (https://tcga-data.nci.nih.gov/tcga/). The metabolic pathway annotation tables were manually curated from the KEGG database, the human-GEM and the fruitly-GEM metabolic atlas^60^.

For a given tumor sample, an enrichment score for a specific metabolic pathway was calculated through single-sample gene-set enrichment analysis (ssGSEA) based on gene expression data. The similarity between a give human tumor sample and the fly *scrib* mutant tumor was then calculated by correlation test using the enrichment scores of 72 conserved metabolic pathways. We set a cutoff on adjusted p-values of correlation coefficients to classify human tumors into metabolically similar and non-similar groups. We then used available somatic mutation information from TCGA and applied Fisher’s exact test to examine significantly-enriched or lost mutation events in human tumors metabolically similar to fly tumors in comparison with human tumors metabolically non-similar to fly tumors. The processed pan-cancer expression data and somatic mutation information of TCGA are provided by UCSC Xena (https://xenabrowser.net/datapages/). To verify our findings using the TCGA dataset, we applied the same methods on the expression and mutation data of glioma samples from CGGA. The gene expression data and somatic mutation information of CGGA are publicly available at CGGA website (https://www.cgga.org.cn/index.jsp).

For bulk RNA-sequencing analysis, the transcript abundance (as denoted by FPKM, Fragments Per Kilobase Per Million reads) of transposons were counted by using piPipes as described in Bo W. Han et al., 2015^61^. Gene expression heatmaps are generated using pheatmap package and PCA analysis was generated using FactoMineR and ggplot2 package. Single cell data merging and gene expression plots were performed using Seurat package^62^.

## Author contributions

Conceptualization, Y.Y., J.G.W.; methodology, Z.C., X.M.Z., M.X.D., C.Y.L., T.T.N., M.L., J.G.W., K.D., T.I.; investigation, Z.C., X.M.Z., M.X.D., C.Y.L., T.T.N.; analysis, Z.C., X.M.Z., M.X.D., C.Y.L., T.T.N.; writing, Y.Y.; funding acquisition, Y.Y., J.G.W., T.I..

## Acknowledgements

We thank the Bloomington Drosophila Stock Center, Vienna Drosophila Resource Center for stocks. We thank Dr. Chris Doe, Dr. Trudi Schupbach, Dr. Rongwen Xi, Dr. Zhao Zhang, and Dr. Robert Campbell for providing fly stocks. This work was supported by grants to Yan Yan from the Research Grants Council of the Hong Kong Special Administrative Region (Grant Number GRF16103620, GRF16104324, T13-602/21N) and from Shenzhen Science and Technology Innovation Commission (Grant Number JCYJ20200109140201722), to Jiguang Wang (Seed fund of the Big Data for Bio-Intelligence Laboratory Z0428 from The Hong Kong University of Science and Technology and Padma Harilela Professorship), to Toyotaka Ishibashi from the National Natural Science Foundation of China (Grant Number 32170548).

