## Supplemental Figure Legend for "Epigenetic reprogramming induced by Acetyl-CoA and SAM depletion is an evolutionarily-ancient path to malignant growth"

**Supplemental Figure 1 The rise of Ldh+ cell population in the *dlg* mutant tumors is regulated by HIF-1 and Pten.**

(A) The *dlg* mutant tumors expressing a Ldh-GFP reporter and a *HIF-1* RNAi construct. Scale bar: 10μm.
(B) The *dlg* mutant tumors expressing a Ldh-GFP reporter and a *Pten* RNAi construct. Scale bar: 10μm.
(C) Quantification of Ldh-GFP+ cell ratio in the *dlg* mutant tumors when *HIF-1* RNAi or *Pten* RNAi is expressed in comparison with control. Control, n=30; *HIF-1* RNAi, n=29; *Pten* RNAi, n=19.

**Supplemental Figure 2: Heatmap plot of the expression of glucose metabolic pathway gene sets in the *scrib* mutant tumors over time.**

**Supplemental Figure 3: Perturbation of AhcyL1 and AhcyL2 does not affect normal wing imaginal disc growth.**

(A) Wing imaginal discs stained for actin (gray) and DNA (blue). Scale bar: 10μm.
(B) Quantification of wing imaginal disc size. For (A-B), control genotype: c855aGal4/+, n=16; Experimental group genotypes: *c855aGal4/AhcyL1^RNAi^, n=19; AhcyL2^RNAi^/+;* *c855aGal4/+, n=16.*

**Supplemental Figure 4: Plot of numbers of human tumor samples by cancer types with similar metabolic signatures to the fly *scrib* mutant tumors.**

**Supplemental Figure 5: Plot of patient survival curves by cancer types comparing human tumors with high metabolic similarity to fly tumors and the non-similar group.**

**Supplemental Figure 6: Analysis of 185 human glioma samples from CGGA database.**

(A) Plot of the distribution of adjusted p-values for metabolic similarity tests between human glioma samples and the fly *scrib* mutant tumors.
(B) 51 out of 185 human glioma samples exhibit metabolic signatures similar to those of the fly *scrib* mutant tumors.
(C) Patients with glioma metabolically similar to the fly tumors are significantly younger than others.
(D) Human glioma samples metabolically similar to the fly tumors acquire a lower number of mutations than other samples.
(E) Human glioma samples metabolically similar to the fly tumors show an enrichment of *IDH1* mutations.
(F) Patients with glioma metabolically similar to the fly tumors have better survival outcomes than the non-similar group.

**Supplemental Table 1 List of metabolite concentrations from the control and the *dlg* mutant larvae as measured in the 600MRM metabolite analysis with LC-MS.**

**Supplemental Table 2 The expression of transposons in the *scrib* mutant tumors over time.**

**Supplemental Table 3 List of conserved metabolic genes and pathways in *Drosophila Melanogaster* and human.**
