## Supplementary figures and images for "Epigenetic reprogramming induced by Acetyl-CoA and SAM depletion is an evolutionarily-ancient path to malignant growth"

### Supplemental Figure 1

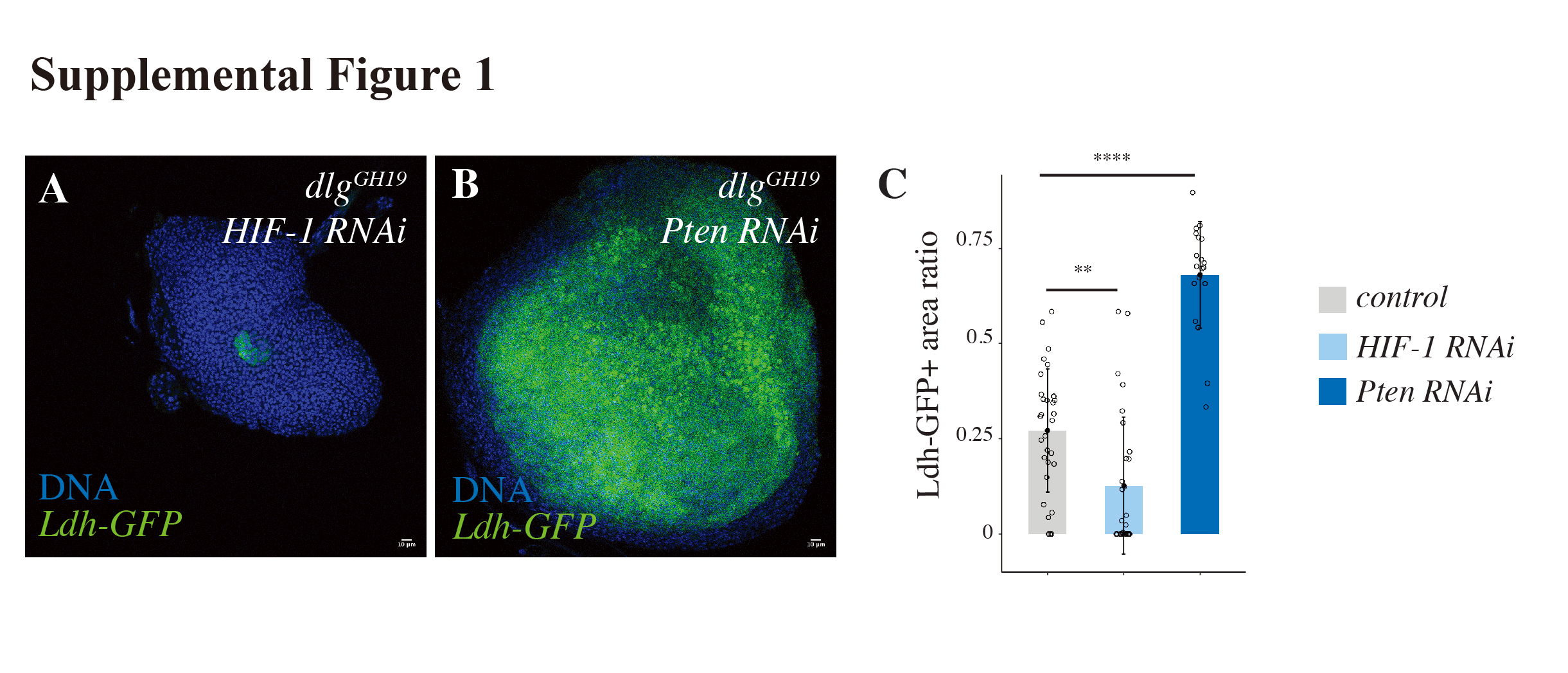

### Supplemental Figure 2

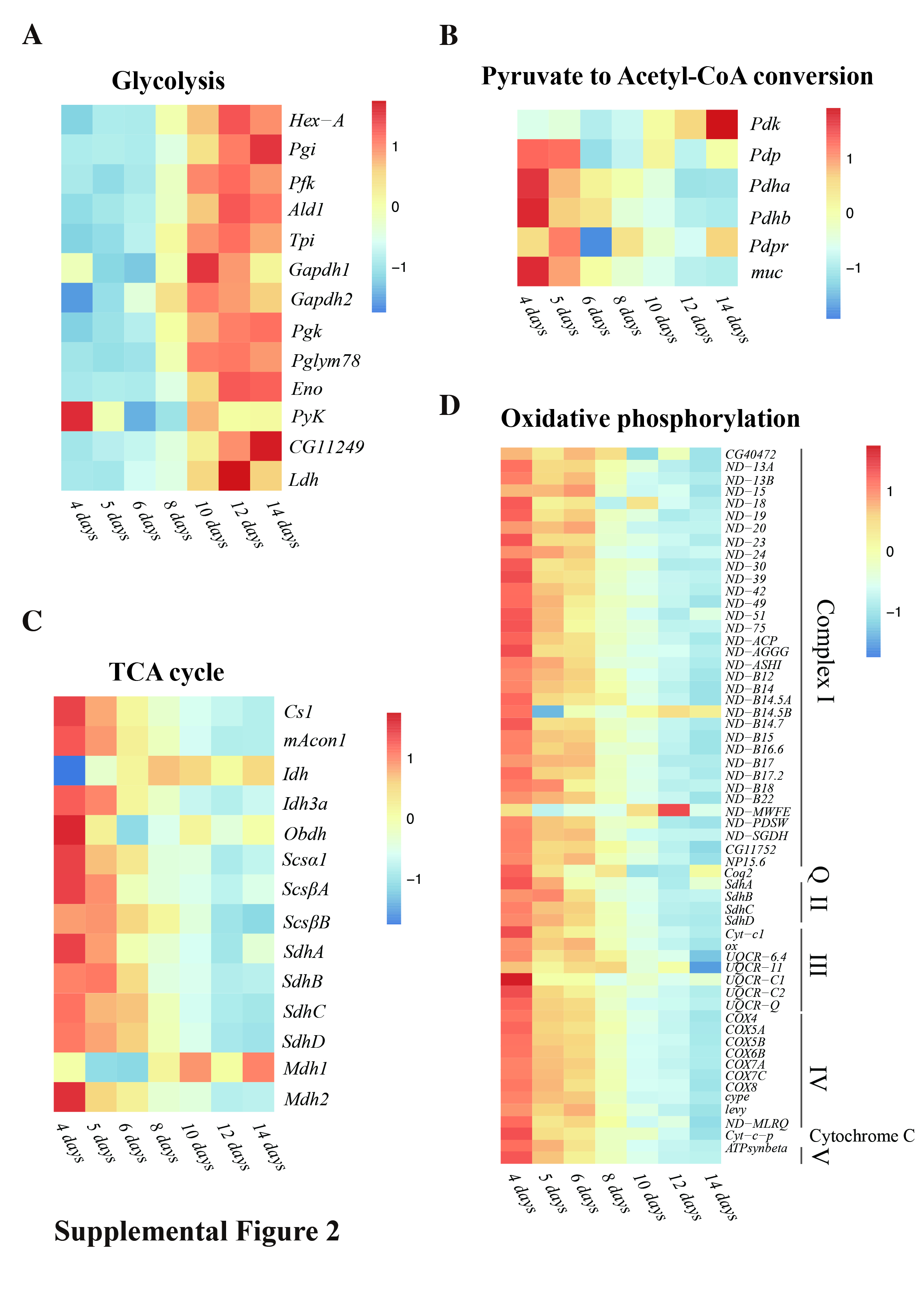

### Supplemental Figure 3

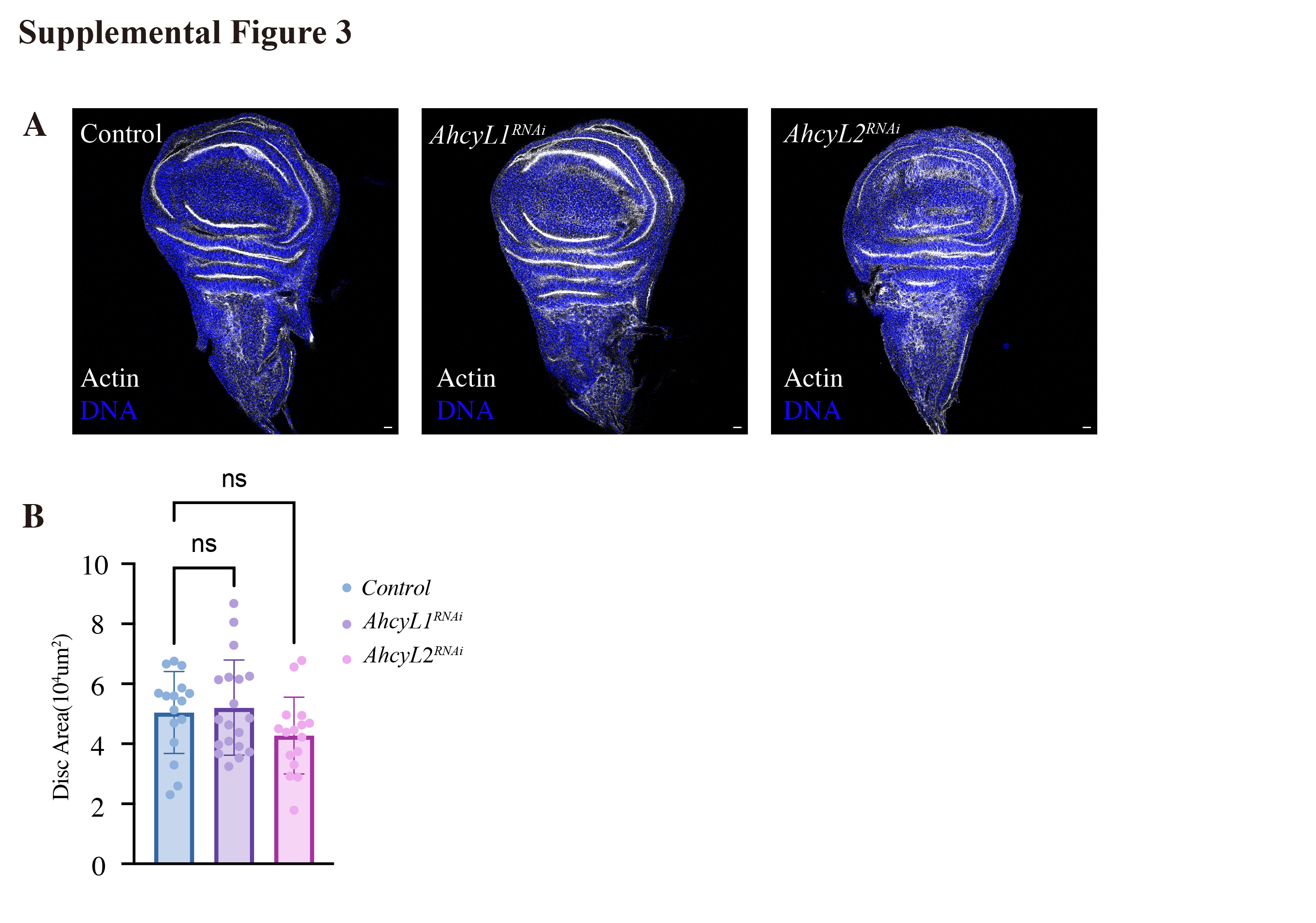

### Supplemental Figure 4

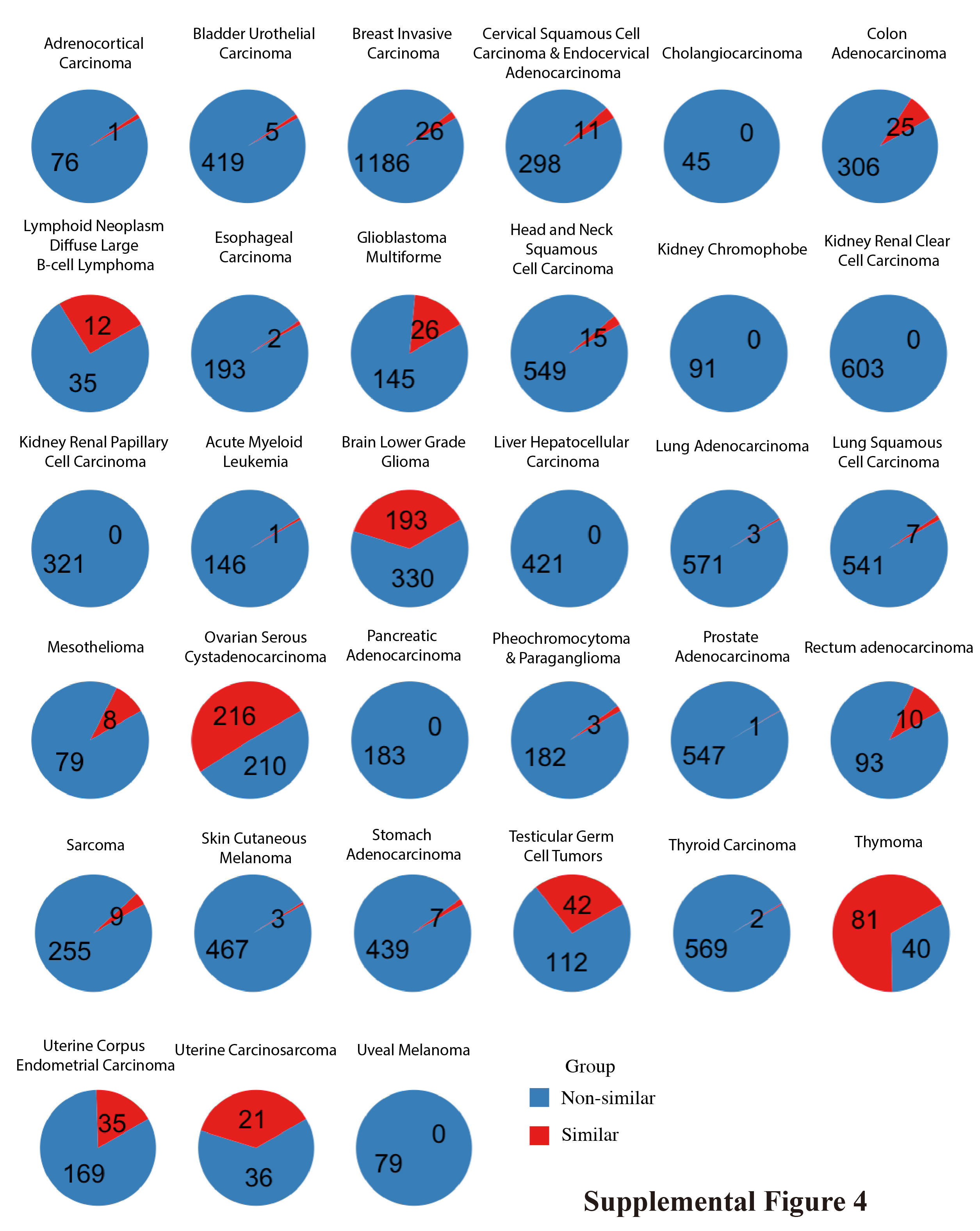

### Supplemental Figure 5

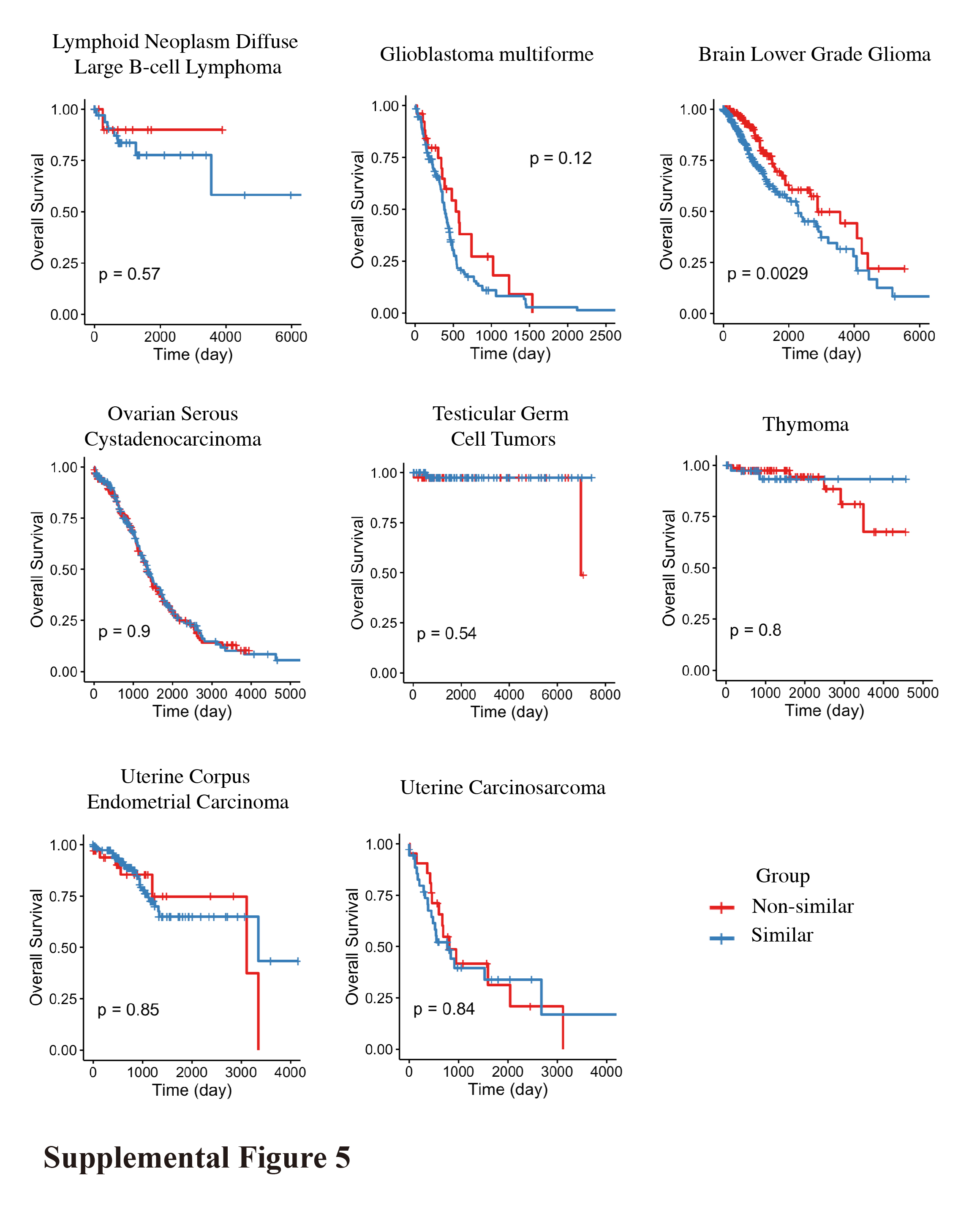

### Supplemental Figure 6

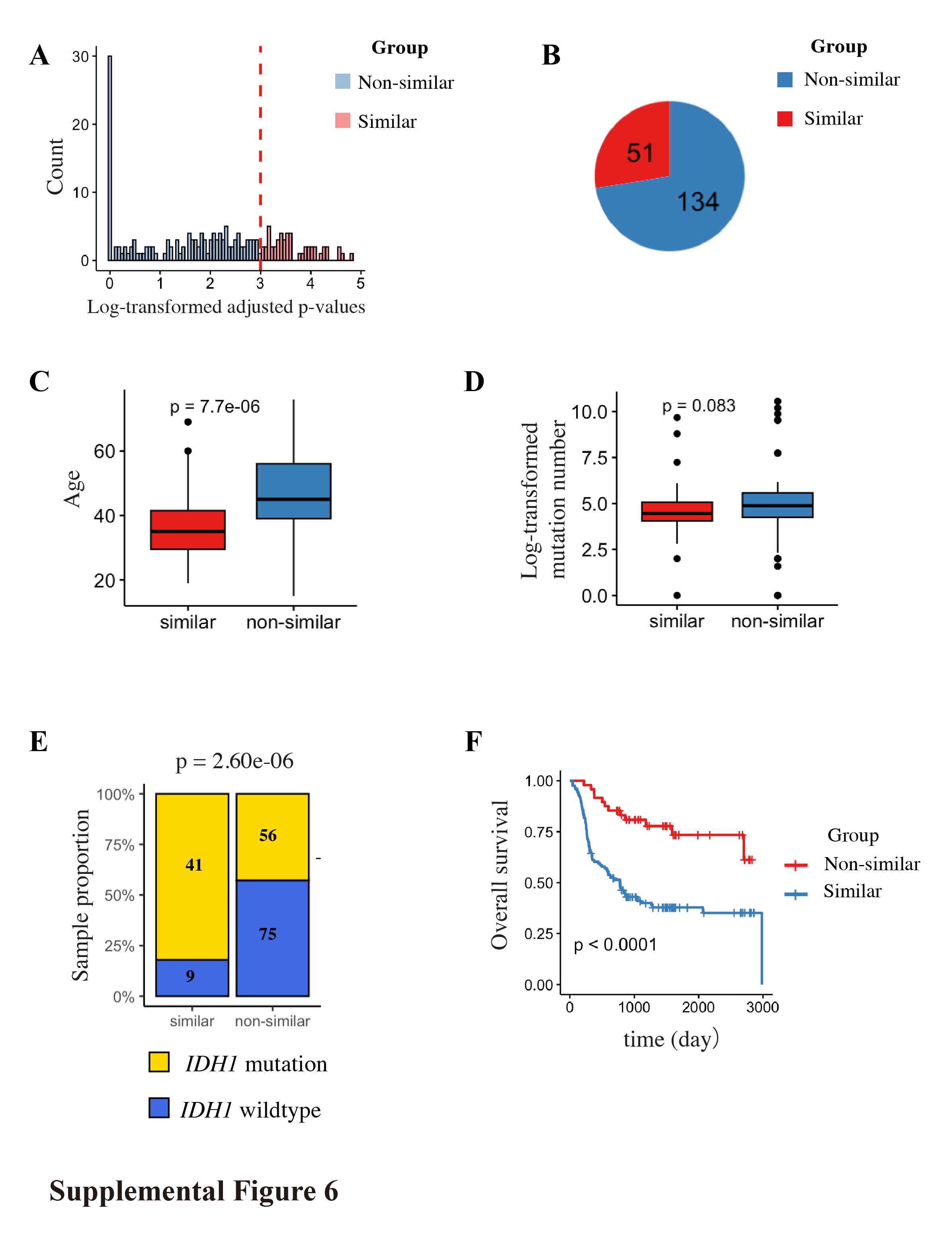
